# Practical and safe method of cryopreservation for clinical application of human adipose-derived mesenchymal stem cells without a programmable freezer or serum

**DOI:** 10.1101/664524

**Authors:** Siqiang Gao, Akiyoshi Takami, Kyosuke Takeshita, Reiko Niwa, Hidefumi Kato, Takayuki Nakayama

## Abstract

**Background:** Adipose-derived mesenchymal stem cells (ADSCs) have emerged as a promising therapeutic modality for cellular therapy because of their rapid proliferation and potent cellular activity compared to conventional bone marrow-derived mesenchymal stem cells (MSCs). Cosmetic lipoaspirates provide an easily obtainable source of ADSCs. Cryopreservation facilitates their clinical application due to increased transportability and pooling of sufficient numbers of cells. However, proper cryopreservation techniques have not been established yet.

**Methods:** We evaluated the post-thaw viability and ADSC functions after cryopreservation with three cryoprotectants (serum containing 10% dimethylsulfoxide (DMSO), serum-free: CP-1^TM^, DMSO-free: SCB-DF^TM^) at two temperature (−80°C, −150°C) and two cell densities: (1 × 10^6^, 7 × 10^6^ cells/mL) for up to 18 months using cryovials. After determining optimal conditions, we also tested if large quantities of ADSCs remained viable after 18 months of cryopreservation in a 100-mL cryobag. Rate-controlled freezing methods or liquid nitrogen storage were not exploited.

**Results:** ADSCs cryopreserved in serum containing 10% DMSO or CP-1^TM^ at −150°C and 7 × 10^6^ cells/mL were most viable (>85%) after 18 months without perturbation of MSC functions. Even suboptimal conditions (−80°C, 1 × 10^6^ cells/mL, no DMSO) assured >80% viability when stored for up to 9 months. Large quantities of ADSCs in a cryobag were properly cryopreserved.

**Conclusions:** A programmable freezer or liquid nitrogen storage is not necessary. CP-1^TM^ is preferable in terms of side effects. Simplified cryopreservation methods (−80°C and no DMSO) can be used for up to 9 months, resulting in reduced infusion toxicities and lower costs.

## Introduction

Mesenchymal stem cells (MSCs) have emerged as a new therapeutic modality for regenerative medicine and immunosuppressive therapy. MSCs can be established from various tissues: bone marrow, adipose tissue, cord blood, dental pulp, umbilical cord, amniotic fluid, etc. Increasing interest has focused on adipose-derived MSCs (ADSCs), because of higher proliferation and cellular activity in addition to the abundant source of adipose tissue obtained by liposuction. MSCs do not elicit allo-reactive lymphocyte responses because they do not express human leukocyte antigen (HLA) class II (1). Thus, third-party ADSCs can be established on a large scale and utilized in many unrelated recipients. Cryopreservation of ADSCs facilitates the clinical application of these cells due to increased transportability and pooling of a sufficient number of cells with the same quality. Compelling studies have shown that strict conditions (stored at an ultra-low temperature, freezing rate controlled by a programmable freezer, usage of cryoprotectant) assure the quality of stem cells, especially hematopoietic stem cells (HSCs), after long-term (more than 10 years) cryopreservation. However, these methods are costly. HSCs should be cryopreserved for a considerably long period because these cells cannot be transplanted into major histocompatibility (MHC) incompatible donors. However, MHC compatibility for MSCs is not necessary as described above, suggesting that more convenient, low-cost methods may be applicable for MSC cryopreservation and subsequent transplantation. Several studies have focused on this aspect of MSC cryopreservation, but most assessed the durability of MSCs in the short term (weeks to months) in small-scale experiments with cryovials (2–4), which are not ideal for clinical settings. Serum containing 10% dimethylsulfoxide (DMSO) has been used for years as a standard cryoprotectant agent (CPA), but this type of CPA has several drawbacks. Serum can provoke an immune reaction and transmit infection. When infused, DMSO exerts side effects such as nausea, emesis, chills, rigors, and cardiovascular events (5–7). Thus, clinical application of ADSCs requires further development of safe, convenient, low-cost cryopreservation methods. Here, we evaluated the post-thaw viability and ADSC functions after cryopreservation with three different CPAs (serum containing 10% DMSO, serum-free, or serum- and DMSO-free) at two different temperatures (−80°C and −150°C) for up to 18 months with the use of cryovials. After setting optimal conditions for storage, we confirmed that large quantities of ADSCs were still viable after cryopreservation in a 100-mL cryobag for 24 months.

## Materials and methods

### Reagents and cells

Heat-inactivated fetal bovine serum (FBS), α-minimal essential medium, and heat-inactivated horse serum were purchased from Gibco-BRL (Invitrogen, Carlsbad, CA). DMSO, hydrocortisone, and 2-mercaptoethanol were from Sigma–Aldrich (St. Louis, MO). The cryoprotectant solutions CP-1™ and Stem-CellBanker^TM^ DMSO FREE GMP Grade (hereafter abbreviated as SCB-DF) were purchased from Kyokuto Pharmaceutical Industrial (Tokyo Japan) and TakaraBio (Shiga, Japan), respectively. For the final freezing medium (hereafter abbreviated as CP-1), 32 mL 25% human serum albumin was added to 68 mL CP-1^TM^ just before use. Human ADSCs were established from adipose tissues to be discarded following skin flap operations after obtaining informed consent (n = 3), as described previously (8).

### Cryopreservation

Human ADSCs were plated onto T75 flasks and cultured until cells reached a enough number for cryopreservation. Prior to freezing, ADSCs (80% confluency) were detached from the flask walls with trypsin/EDTA, and viable cells were counted with the trypan blue exclusion method. After centrifugation, supernatant was removed completely, and cells were resuspended in one of three types of freezing medium: 10% DMSO in FBS (hereafter abbreviated as DMSO), CP-1, or SCB-DF, at two different cell densities: 1 × 10^6^ or 7 × 10^6^ cells/mL in 0.2-mL cryopreservation vials (0.1 mL/vial), and then stored in a −80°C deep freezer or a −150°C ultra-low temperature freezer for 3, 9, or 18 months. In another setting, cell suspensions of ADSCs were prepared at approximately 1 × 10^6^ cells/mL in a total volume of 50 mL and then placed in 100-mL cryobags (Terumo, Tokyo, Japan). The cryobags containing ADSCs were directly placed into a −150°C ultra-deep freezer and stored for 24 months.

### Cell thawing and cell recovery rate

Frozen cryovials or cryobags were quickly thawed in a circulating 37°C water bath by gently shaking and agitating until the ice mass disappeared. Then, the cell suspension from cryovials was washed with culture medium once, and recovery rates were evaluated with the trypan blue exclusion method. The recovery rates were calculated as follows: (viable cell number/total cell number) × 100 (%). The rest of the cells were grown in culture to evaluate hematopoietic-supporting activities, surface antigen profiles, and differentiation abilities into adipocytes and osteoblasts.

### Fluorescence-activated cell sorting (FACS) analysis and differentiation assay

After the freeze-thawing process, ADSCs were incubated with the following fluorescently labeled mouse anti-human monoclonal antibodies: CD14-FITC (Immunotech, Marseille, France), CD29-PE (BD Biosciences, San Jose, CA), CD34-PerCP (BD Biosciences), CD45-FITC (Immunotech), CD73-PE (BD Biosciences), CD90-PE (eBioScience, Pittsburgh, PA), CD105-PE (eBioScience), HLA Class1-PE (eBioScience), and HLA-DR-FITC (Immunotech). Data were acquired on a MoFloTM XDP cell sorter (Beckman Coulter, Brea, CA) by collecting a minimum of 10,000 events and analyzed with Summit software V5.3.0.10325 (Beckman Coulter). Background fluorescence was assessed by staining with isotype-matched antibodies. The multilineage potential of ADSCs was confirmed as described elsewhere (9). Briefly, freshly established ADSCs and ADSCs after the freeze-thawing process were exposed to adipogenic formulas (R&D Systems, Minneapolis, MN) for 14 days or to osteogenic formulas (R&D Systems) for 21 days. Accumulation of intracellular lipid-rich vacuoles resulting from adipogenic differentiation was assessed by Sudan III staining. Osteogenic differentiation was specifically evaluated by von Kossa staining to detect calcium deposition.

### Coculture of human CD34-positive progenitor cells with human ADSCs

Coculture of human CD34-positive peripheral blood stem cells and human ADSCs was performed as described previously (10). Briefly, feeder layers comprised of intact ADSCs or ADSCs after the freeze-thawing process were pre-established (80% confluency) in 48-well plates pre-coated with gelatin. Then, CD34-positive cells (1.0 × 10^4^ cells per well), purchased from the RIKEN BRC through the National Bio-Resource Project of the MEXT, were suspended in long-term culture medium (α-MEM containing 12.5% horse serum, 12.5% FBS, 1 μM hydrocortisone, and 50 μM 2-mercaptoethanol) and applied to the feeder layers. The cocultures were incubated for 4 weeks with replenishment of long-term culture medium twice a week, and proliferation of floating cells was assessed in three randomly selected microscopic fields (200×). At the end of the coculture, the progenitor content of each well was assayed with methylcellulose colony assays as described previously (11). All adherent cells in each well were detached by trypsinization and plated in 3 mL Methocult^TM^ H4034 (Stemcell Technologies, Vancouver, Canada). After 14 to 16 days of incubation, erythroid (burst forming unit-erythroid [BFU-E]), myeloid (colony forming unit-granulocyte, macrophage [CFU-GM]), colony forming unit-granulocyte [CFU-G], colony forming unit-macrophage [CFU-M]), and multipotent (colony forming unit-granulopoietic, erythroid, macrophage, megakaryocyte [CFU-GEMM)]) progenitor cells were quantified under a microscope. Experiments were repeated two times.

### Statistical analysis

Results were expressed as the mean ± standard deviation (SD). The statistical significance of group differences was evaluated with the Student’s *t* test using Excel software (Microsoft, Redmond, WA). All statistical tests were two-sided.

## Results

### Influence of storage conditions on the post-thaw recovery rate of human ADSCs

To identify the optimal storage conditions for human ADSCs, cells were suspended in three different CPAs: DMSO (open columns), CP-1 (black columns), or SCB-DF (gray columns), at two different cell densities: 1 × 10^6^ or 7 × 10^6^ cells/mL in 0.2-mL cryopreservation vials (0.1 mL/vial), and then stored in a −80°C deep freezer or a −150°C ultra-low temperature freezer for 3, 9, or 18 months. The stored cells were rapidly thawed in a water bath at 37°C, and viable cells were counted using the trypan blue exclusion method under a microscope. Cells in all subgroups showed minimal cell aggregation.

When stored at −80°C (Figure 1A), the post-thaw recovery rate of human ADSCs was stable after 9 months, but rapidly and significantly decreased after 18 months. None of the CPAs or cell densities tested prevented the decrease in the post-thaw recovery rate at 18 months. However, DMSO, CP-1, and storage at high cell density (7 × 10^6^ cells/mL) were superior to SCB-DF and storage at low cell density (1 × 10^6^ cells/mL).

**Figure 1.**
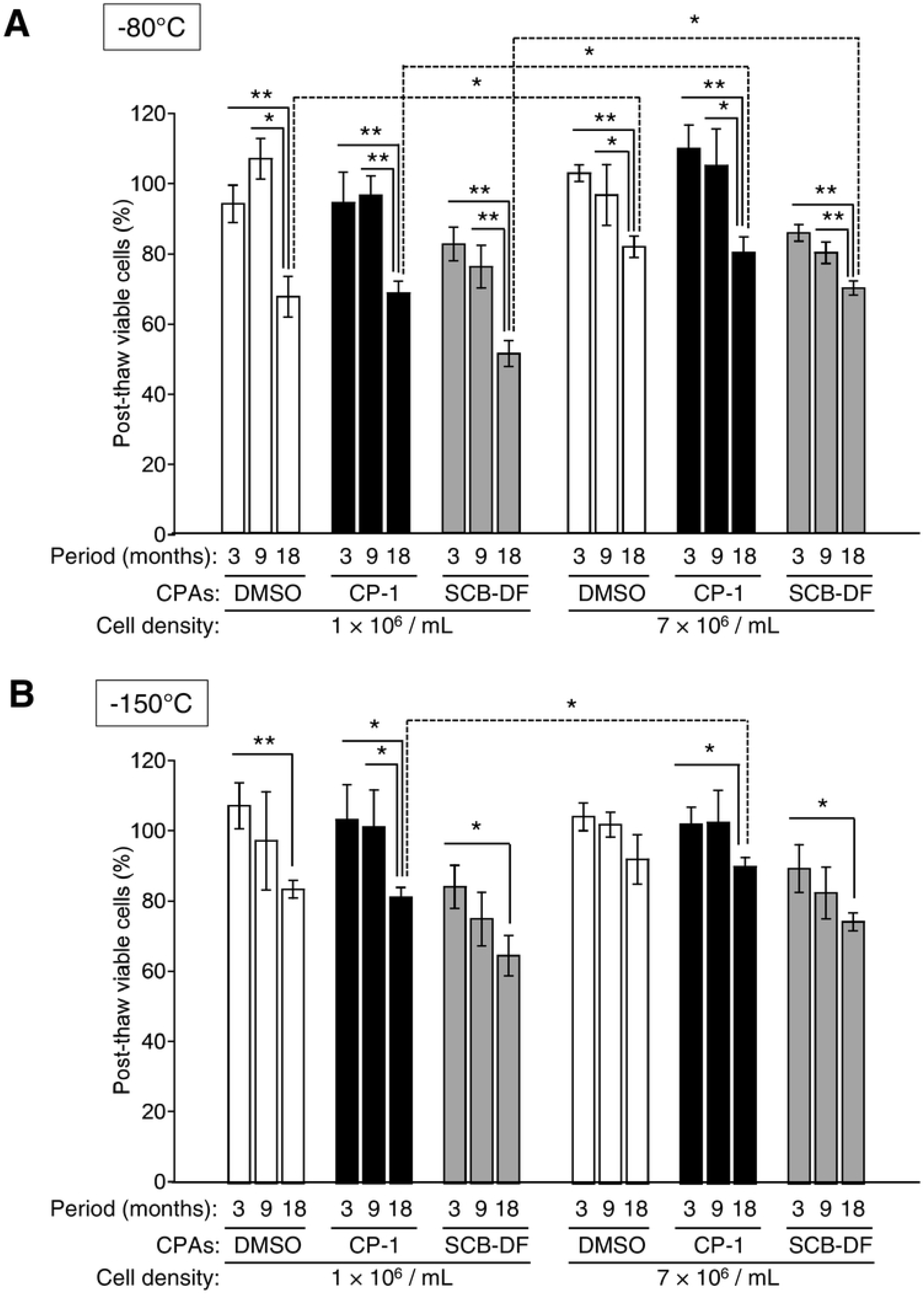
Post-thaw recovery rate of human ADSCs in various freezing conditions. Prior to cryopreservation, human ADSCs were suspended in three different CPAs: DMSO (white columns), CP-1 (black columns), or SCB-DF (gray columns), at two different cell densities: 1 × 10^6^ or 7 × 10^6^ cells/mL in 0.2-mL cryopreservation vials (0.1 mL/vial), and then stored in a −80°C deep freezer (A) or a −150°C ultra-low temperature freezer (B) for 3, 9, or 18 months. The stored cells were rapidly thawed in a water bath at 37°C, and viable cells were counted with the trypan blue exclusion method. The percent post-thaw viable cells was calculated as follows: viable cells after thawing (cells/mL)/viable cells before cryopreservation (cells/mL) × 100 (%). The results shown reflect the mean (± SD) of triplicate determinations. The asterisks denote statistical significance (**P < 0.01; *P < 0.05).

When stored at −150°C (Figure 1B), the post-thaw recovery rate of human ADSCs decreased over time, similar to storage at −80°C. However, cell viability was better in all subgroups at 18 months compared to storage at −80°C. Additionally, storage at the high cell density (7 × 10^6^ cells/mL) maintained greater viability of ADSCs than storage at the low cell density (1 × 10^6^ cells/mL) in each CPA. Minimal cell damage was observed in cells stored at high cell density (7 × 10^6^ cells/mL) with DMSO or CP-1, even after 18 months.

These data suggest that optimal storage conditions for human ADSCs are cryopreservation in DMSO or CP-1, low temperature (−150°C), and high cell density (7 × 10^6^ cells/mL). ADSCs can be safely cryopreserved for at least 18 months in such a condition.

### The hematopoiesis-supporting ability of human ADSCs is not affected by long-term cryopreservation

To assess the quality of human ADSCs after long-term cryopreservation, we focused on the hematopoiesis-supporting ability of ADSCs, because ADSCs support human CD34^+^ HSCs to a greater degree than conventional bone marrow-derived MSCs. In cocultures, we observed that CD34^+^ cord blood cells overlaid onto fresh ADSCs (hatched column) or ADSCs cryopreserved in CPAs (DMSO: open columns, CP-1: black columns, SCB-DF: gray columns) formed clusters of round cells that were unattached or tightly attached to the monolayers (Figure 2A, left upper panel), which was consistent with a previous report (10). We compared the number of these nonattached cells at the indicated time points, and found that the number of nonadherent cells was not significantly different among the four subgroups (Figure 2A, right upper panel). After 28 days of coculturing, adherent cells were quantitated with colony-forming cell assays. The cocultured cells formed CFU-GEMM, CFU-G, and CFU-M in the four subgroups. The major lineages of the colonies were CFU-G and CFU-GM (Figure 2B), consistent with our previous report (12). We found no significant difference in the proportion of each lineage or the total number of colonies among the four subgroups (Figure 2B).

**Figure 2.**
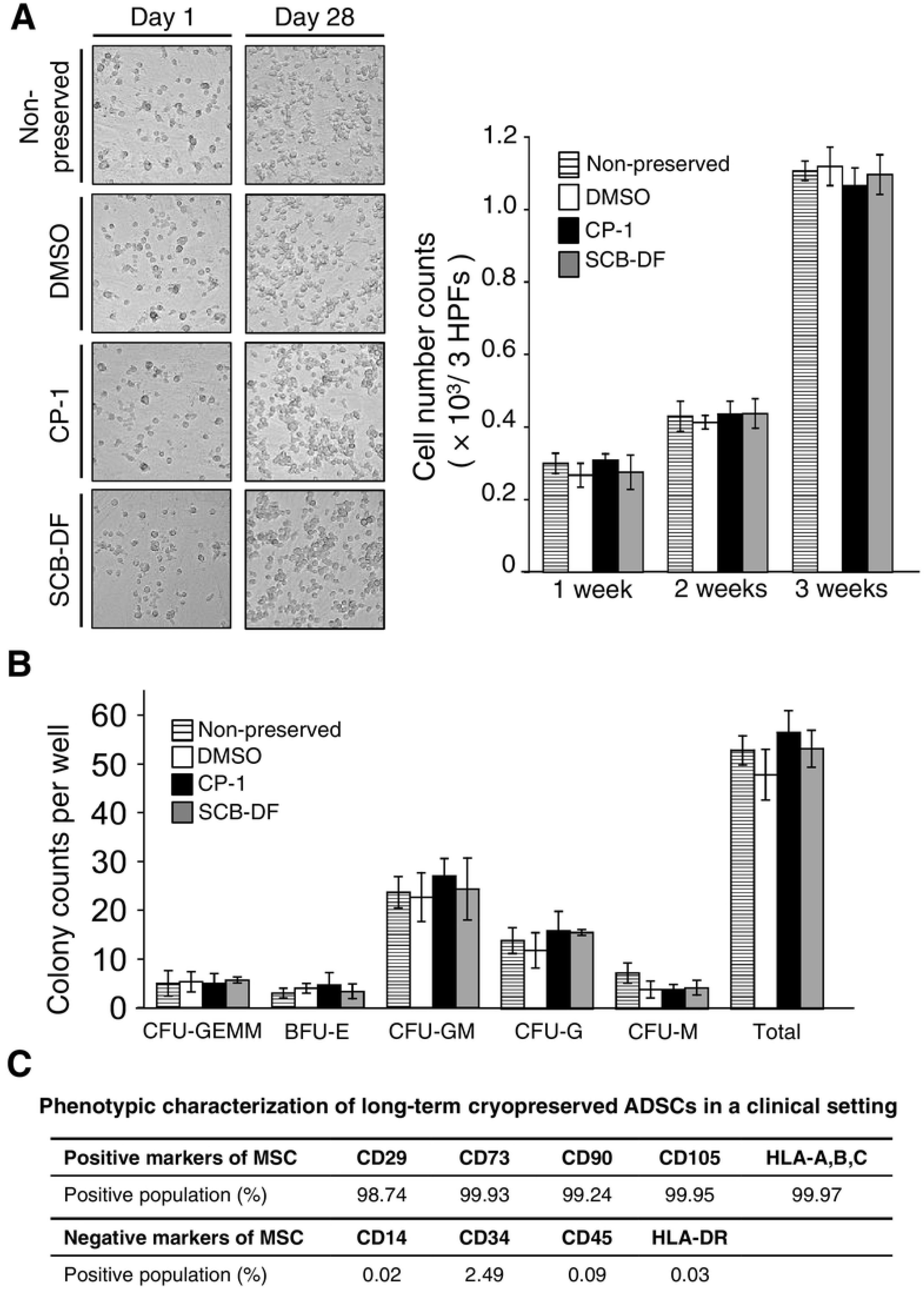
Quality assessment of human ADSCs after long-term cryopreservation. Hematopoiesis-supporting ability and CD marker expression. (A) Cells treated with DMSO or CP-1 were more viable than those treated with SCB-DF. The quality of ADSCs treated with DMSO, CP-1, or SCB-DF was analyzed by examining the hematopoiesis-supporting ability of human ADSCs. Human ADSCs cryopreserved at a density of 7 × 10^6^ cells/mL in 0.2-mL cryovials at −150°C for 18 months with DMSO (white columns), CP-1 (black columns), or SCB-DF (gray columns) were thawed, cultured, and then used for coculture assays with CD34^+^ HSCs from cord blood. Non-preserved human ADSCs (lined columns) were used as a control. Cell layers of human ADSCs were established on 0.5% gelatin pre-coated 48-well plates (80% confluency). CD34^+^ HSCs were applied on the feeder cell layers (1.0 × 10^4^ cells/400 μL/well, three wells per subgroup), and cultured at 37°C for 4 weeks with refreshment of culture medium twice a week. Nonadherent viable cells were photographed (left panel) and counted (three high-power fields were randomly selected) at the indicated time points (right panel). Adherent cells were quantitated with colony-forming cell assays. (B) After 28 days of coculturing, all adherent cells were detached by trypsinization. Then, viable cells were counted with the trypan blue exclusion method and plated in 3.3 mL methylcellulose (2 × 10^4^ cells). The plates were incubated for 14-16 days, after which progenitors were scored (lower panel). The results represent the mean (±SD) of three independent experiments. (C) Long-term effects of cryopreservation on human ADSC phenotype in a clinical setting. Human ADSCs, which were cryopreserved in 100-mL cryobags for clinical use in CP-1 (approximately 1 × 10^6^ cells/mL in a total volume of 50 mL) at −150°C for 24 months, were quickly thawed in a water bath, and CD markers were evaluated with FACS. Representative results of three independent experiments are shown.

### Long-term cryopreserved human ADSCs in a clinical setting retained surface antigen profiles specific to human MSCs and differentiation potential

Large quantities of cells for infusion are required for clinical treatment. Thus, we next cryopreserved human ADSCs in 100-mL cryobags for clinical use for 24 months in optimal conditions: in CP-1 (approximately 1 × 10^6^ cells/mL in a total volume of 50 mL) at −150°C. After 24 months of cryopreservation, cells were quickly thawed in a water bath at 37°C. The viability of cells was 91.9 ± 3.9% (mean ± SD) with minimal cell aggregates. The cells showed no morphological changes and propagated well, similar to fresh ADSCs. FACS analysis showed that post-thawed human ADSCs were positive for CD29, CD73, CD105, CD90, and HLA-ABC, and negative for CD45, CD34, CD14, and HLA-DR (Figure 2C). These phenotypes met the minimal criteria of MSCs as defined by the International Society for Cell Therapy (13). To confirm the differentiation ability of the frozen and thawed human ADSCs, cells were exposed to adipogenic formulas for 14 days or osteogenic formulas for 21 days. Accumulation of intracellular lipid-rich vacuoles resulting from adipogenic differentiation was assessed with Sudan III staining. Osteogenic differentiation was specifically evaluated with von Kossa staining to detect calcium deposition. Lipid drops were clearly detected inside cells, but calcium deposition was minimally observed (data not shown).

## Discussion

In this report, we evaluated the post-thaw viability and ADSC functions after cryopreservation with three different cryoprotectants (serum containing 10% DMSO, serum-free, or DMSO- and serum-free), at two different cell densities: 1 × 10^6^ or 7 × 10^6^ cells/mL, at two different temperatures (−80°C and −150°C) for up to 18 months. We did not exploit rate-controlled freezing methods or liquid nitrogen storage. We found that the following conditions were optimal: higher cell density (7 × 10^6^ cells/mL), usage of DMSO, and lower storage temperature (−150°C) (Figure 1A and 1B). However, even suboptimal conditions (−80°C, 1 × 10^6^ cells/mL, no DMSO) in which cells were stored up to 9 months produced a post-thaw recovery rate >80% (Figure 1A and 1B). These facts suggest that rate-controlled freezing methods or liquid nitrogen storage are not necessary for cryopreservation of ADSCs over a short-to mid-term period. Serum containing 10% DMSO has been used for years as a standard cryoprotectant agent, because DMSO can penetrate into cells and prevent the formation of intracellular ice crystals and subsequent membrane rupture. Our current experiments showed that post-thaw cell recovery was significantly low in ADSCs cryopreserved with SCB-DF (Figure 1A and 1B), suggesting an important role for DMSO. However, DMSO, when infused into the body, exerts side effects such as nausea, emesis, chills, rigors, and cardiovascular events in a dose-dependent manner (5–7). CP-1 is a 25% human serum albumin solution that is mixed just before use and added to an equal volume of cell suspension, yielding a final freezing medium with 5% DMSO, which has low risk for side effects. In Japan, CP-1 has been clinically used for cryopreservation of HSCs for decades with few side effects reported (14, 15). Thus, CP-1 may be a safe, easy-to-prepare, low-cost cryopreservation medium for ADSCs.

Freezing cells and thawing cryopreserved cells properly is crucial to the viability and functionality of the cells. Freezing and thawing a large number of cells, which is imperative for clinical application, in cryobags takes longer compared to cryovials. However, the viability of human ADSCs cryopreserved in cryobags at −150°C for 24 months was over 90% with minimal cell clumps. The morphology, growth rate, surface antigen profiles, and adipogenic differentiation, but not osteogenic differentiation, of these cells were the same as those of fresh ADSCs. The osteogenic potential of ADSCs is inferior to that of BMSCs (16), and the cryopreservation process decreases the osteogenic differentiation ability of ADSCs (17, 18). Little was known about how long-term cryopreservation affects the hematopoietic-supporting ability of ADSCs, one of the major features of ADSCs (19). Here, we show that both cryopreserved and fresh ADSCs generated almost equal numbers of granulocytes and progenitor cells from human HSCs using *in vitro* coculture and progenitor assays (Figure 2B), even though the cell recovery rates of ADSCs stored with SCB-DF were lower compared with those with DMSO or CP-1 (Figure 1A and 1B). These facts clearly suggest that cryopreservation can perturb the ability of osteogenic differentiation of ADSCs but not the hematopoietic-supporting ability.

Together, these observations provide helpful information about safe, cost-effective cryopreservation of ADSCs that could be used in the clinical setting.

## Acknowledgements

The authors wish to thank Ms. Yukiji Ando and Ms. Rie Goto for their contributions to various aspects of this work.

